# Methodological considerations in analyzing synchronization of resting-state brain networks with the intrinsic electrical rhythm of the stomach: Advantages of weighted phase-locking

**DOI:** 10.1101/2021.09.20.461120

**Authors:** Ann S. Choe, Bohao Tang, Kimberly R. Smith, Hamed Honari, Martin A. Lindquist, Brian S. Caffo, James J. Pekar

## Abstract

**Purpose:** To evaluate the amplitude-weighted phase-locking value (awPLV) as a measure of synchronization of brain resting-state networks (RSNs) with the gastric basal electrical rhythm (BER).

**Methods:** A recent study combined rsfMRI with concurrent cutaneous electrogastrography (EGG), in a highly-sampled individual who underwent 22 scanning sessions (two 15-minute runs per session) at 3.0 Tesla. After excluding three sessions due to weak EGG signals, 9.5 hours of data remained, from which 18 RSNs were estimated using spatial independent component analysis. Previously, using the phase-locking value (PLV), three of the 18 RSNs were determined to be synchronized with the BER. However, RSN power fluctuations in the gastric frequency band could reduce sensitivity of PLV. Accordingly, the current reanalysis used awPLV to unweight contributions from low power epochs. Mismatched EGG and rsfMRI data (from different days) served as surrogate data; for each RSN, empirical awPLV was compared with chance-level awPLV using a Wilcoxon rank test. P-values were adjusted using with a false discovery rate of 0.05. Additionally, simulations were performed to compare PLV and awPLV error rates under settings with a known ground truth.

**Results:** Simulations show high false-negative rates when using PLV, but not awPLV. Reanalysis of the highly-sampled individual data using awPLV indicates that 11 of the 18 RSNs were synchronized with the BER.

**Conclusion:** Simulations indicate that awPLV is a more sensitive measure of stomach/brain synchronization than PLV. Reanalysis results imply communication between the enteric nervous system and brain circuits not typically considered responsive to gastric state or function.

## INTRODUCTION

This report introduces weighted phase-locking as a sensitive measure of the synchronization of restingstate BOLD fMRI signals with the intrinsic electrical rhythm of the stomach. The gastric basal electrical rhythm (BER) is generated continuously and intrinsically by a network of myenteric interstitial cells of Cajal (1, 2), which creates and sustains a traveling electrical slow wave that governs peristalsis when the stomach contains food or chyme. The *ca*. 0.05 Hz gastric BER is transmitted to the brain chiefly by vagal afferents. Recently, we combined concurrent cutaneous electrogastrography (EGG) with resting-state fMRI (rsfMRI), in a highly-sampled individual experimental design. Eighteen resting state networks (RSNs) were estimated using spatial independent component analysis (ICA). Using the phase-locking value (PLV), three of those 18 RSNs were found to be phase-locked with the gastric basal electrical rhythm (3). However, RSN time courses did not consistently sustain strong signal in the gastric frequency band; this suggests, consistent with a report on assessing synchronization in electrophysiological recordings (4), that noisy phase estimates during epochs of low signal-to-noise reduce the sensitivity of PLV. To address this, the present contribution is a *reanalysis* of those data, using the *amplitude weighted* Phase-Locking Value (awPLV), a measure introduced by Kovach (4) in order to improve estimation of phase synchronization (5, 6). Using awPLV, we now find a *majority* of resting-state brain networks significantly synchronized with the intrinsic rhythm of the stomach. Additionally, we conducted plasmode / Monte Carlo simulations in order to characterize error rates of significance tests based on the PLV and awPLV measures.

## METHODS

### Empirical Methods

As described previously (3), an individual participant gave informed consent to participate in a study approved by the Johns Hopkins Medicine Institutional Review Board. This healthy male volunteer, age 58, with a body mass index (BMI) of 26, underwent 22 scanning sessions at 3.0 Tesla (Philips dStream Ingenia Elition scanner), each at 9 AM (the subject breakfasted at 5:30 AM), over a span of seven weeks. In each session, two 15-minute runs of concurrent EGG and rsfMRI data were acquired; the participant remained in the scanner between runs. A multi-slice SENSE-EPI pulse sequence was used for rsfMRI. The gastric rhythm was monitored using MRI-compatible EGG equipment (BIOPAC MP160 system; BIOPAC Systems Inc, USA). Detailed MRI and EGG methods, including session dates, may be found in our prior report (3). EGG data from three sessions had weak gastric signals and were excluded; the other 19 sessions yielded a total of 9.5 hours of data.

The data processing pipeline is outlined in Figure 1. As reported previously, the rsfMRI data were analyzed using group independent component analysis (ICA) (7–9); resting-state network (RSN) time courses were estimated for each run (3). EGG data were preprocessed as described previously (3); data from the EGG channel with the highest gastric spectral power were bandpass filtered to isolate the signal related to the gastric rhythm. To estimate stomach/brain phase synchronization, the RSN time courses were filtered using the same bandpass filter as the EGG data; the Hilbert transform was used to estimate the analytic phase of each signal; then, for each run, the awPLV was computed between the gastric and (filtered) RSN time courses.

**Fig 1.**
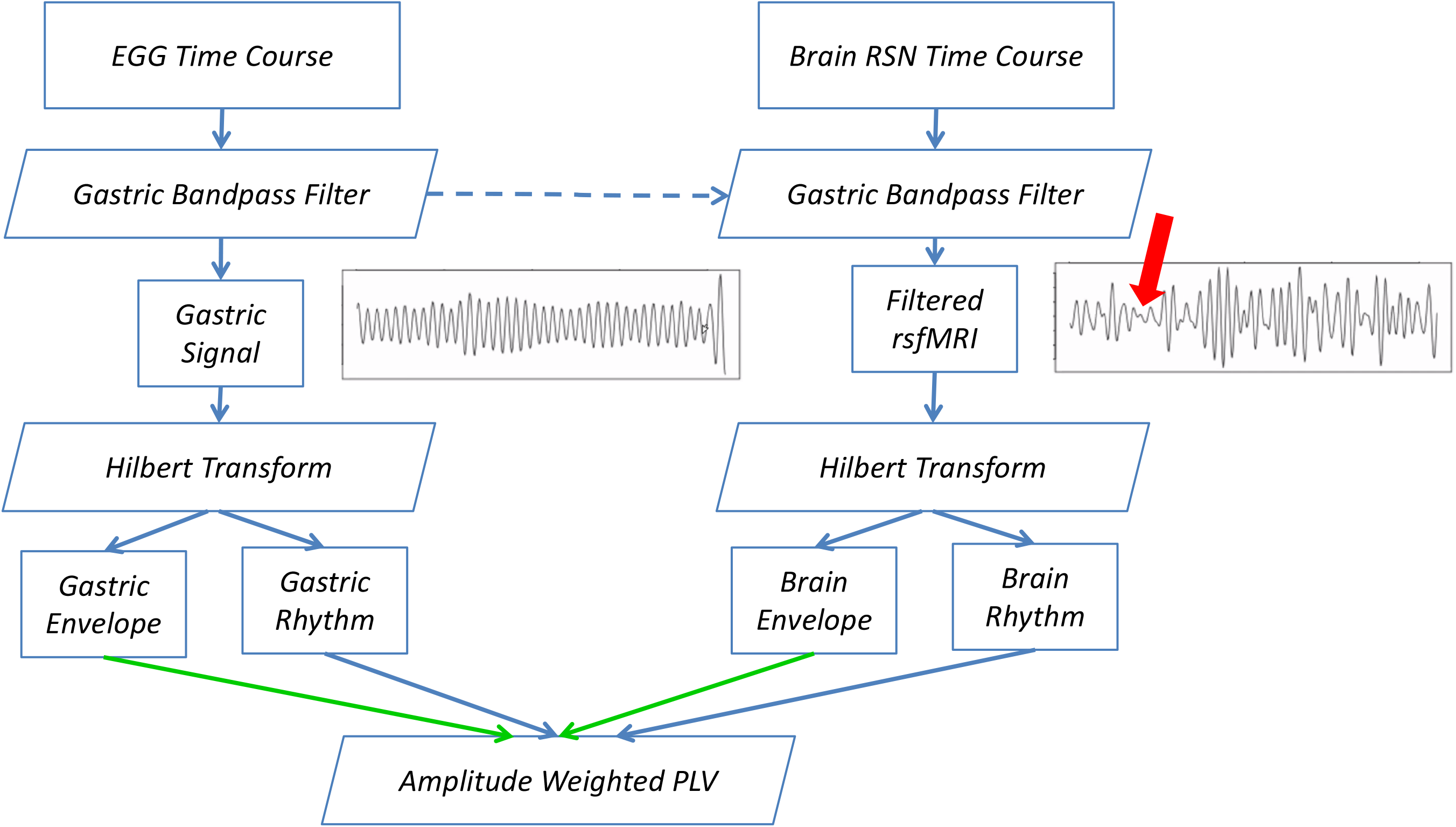
Data analysis pipeline. Plots show empirical data. Broad red arrow indicates a period of low signal that will have noisy (poorly-defined) phase. Green arrows indicate signal amplitudes, which are used for the awPLV measure, but ignored in the PLV measure.

The awPLV between two signals *x*(*t*) and *y*(*t*) is defined as (4):

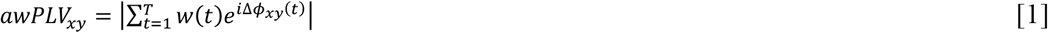

Where the phase difference is Δ*ϕ_xy_*(*t*) = *ϕ_x_*(*t*) – *ϕ_y_*(*t*) and the weights are given in terms of the signal envelope amplitudes:

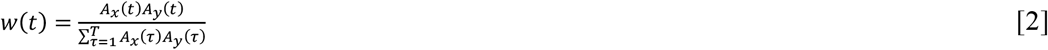

The difference between awPLV and PLV lies in the weighting term. PLV weighs all of the data equally (i.e., w(t) = 1 for all t). Hence, it runs the risk of over-weighing data where the signal amplitude approaches zero (which leads to poor estimation of instantaneous phase at these instances), such as indicated by the broad red arrow in Figure 1. The awPLV addresses this by down-weighing these contributions using the envelope amplitudes, as indicated by the green arrows in Figure 1. As this is a situation that occurs in the RSN time courses, this suggests that awPLV may be a more appropriate metric in this setting.

In order to assess statistical significance, following the approach used earlier for PLV (3), for each RSN, awPLV was calculated for all pairs of “mismatched” data (EGG and rsfMRI data acquired on different days) in order to generate a null distribution of chance-level gastric/brain synchronization. We then tested whether, for each RSN, empirical awPLV differed from chance-level awPLV, using a Wilcoxon rank test. Finally, to correct for multiple comparisons, the p-values for the 18 RSNs were adjusted using the Benjamini-Hochberg procedure with a false discovery rate (FDR) of 0.05, to determine whether RSNs were significantly synchronized with the gastric rhythm.

### Simulation Methods

To understand and contextualize the differing results of statistical inference using the awPLV vs. PLV analyses, we conducted plasmode Monte Carlo simulations. The dual goals of the simulations were to assess the false negative rates of inference based on each of the two metrics, using simulated data with known “ground truth” stomach/brain phase synchronization; and to assess the false-positive rates of inference based on each of the two metrics, using different simulated data, without ground truth stomach/brain phase synchronization. The simulation pipeline is outlined in Figure 2.

**Fig 2.**
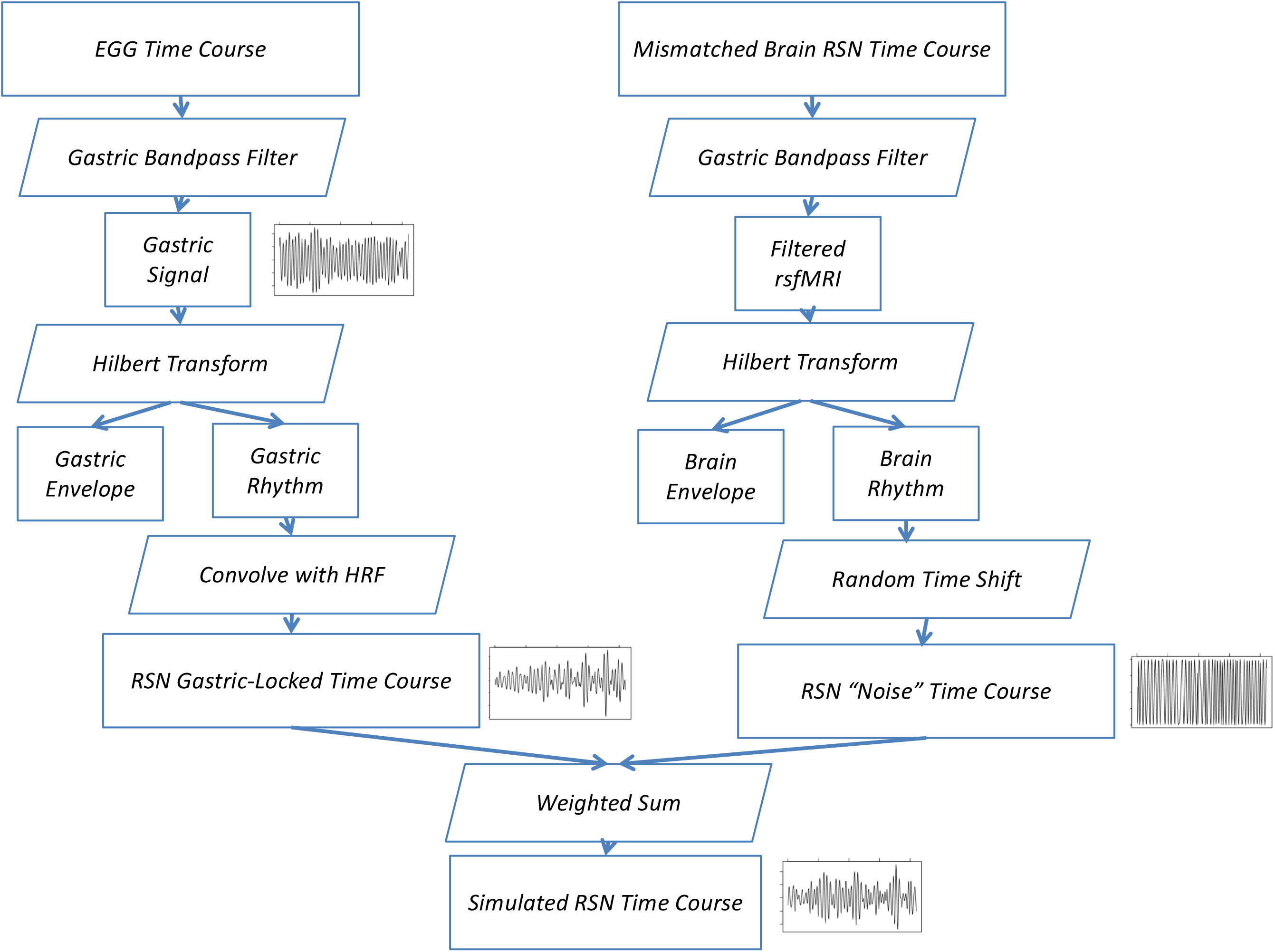
Simulation pipeline. Plots show empirical data (EGG timecourse) and simulated results (others)

To generate simulated brain RSN data with a fixed ground truth gastric synchronization, we took EGG time course data, temporally filtered and Hilbert transformed, as described above, then convolved the resulting gastric rhythm with an estimate of the brain hemodynamic response function to yield a simulated RSN time course with fixed and known EGG phase-locking. To obtain realistic RSN noise to add to these ideal data, we took RSN time course data from a different day, and subjected it to passband temporal filtering and Hilbert transformation, followed by a random time shift. The random time was performed as y(t)=x(t+k) where x(t) is the original signal at time t, k is the random shift and y is the resulting noise. The shift, k, was chosen as uniform on the time range, excluding the first and last 30 seconds and x(t+k) was wrapped around (i.e. t+k was taken modulo the largest time point). The simulated RSN signals were then obtained by a weighted sum of the fixed ground truth signal and the simulated random noise. The weights were chosen to yield a contrast to noise ratio (CNR) level of 20% (20% of the simulated signal is ground truth and 80% of the signal is noise) equivalent to a signal to noise ratio (SNR) of 4. These values were chosen as consistent with our observed data. Simulations were also performed at a CNR level of 30%, to investigate how improved SNR would influence error rate performance. Representative simulated time courses are shown in Figure 2. These simulated ground truth noisy data were used to estimate the rate of missed phase synchronization, or the False Negative Rate (FNR).

To generate simulated null distribution rsfMRI data lacking stomach/brain synchronization, we used the RSN noise as described above. That is, only the noise was used (a CNR of 0). Then, these null simulated data were used to assess the rate of false positives, or the False Positive Rate (FPR).

For each simulation, 2,000 runs were performed, sampling data as described above, from each RSN time course, from each acquired scan.

## RESULTS

### Simulation Results

Simulation results are summarized in Table 1. False Positive Rates from null data were below the specified 0.05 alpha level. False Negative Rates at the empirical CNR level of 20% (corresponding to an SNR of 4) were approximately 77%, decreasing to 18% with an increase in the CNR to 30%.

**Table 1.** Simulation results: False-positive rates (FPR) for null data, and false-negative rates (FNR) at contrast-to-noise (CNR) levels of 20% and 30% (see text), using the PLV and awPLV measures. Mean values are followed by 95% confidence Intervals.

| Error Rate | Phase Synchronization Measure |  |
| --- | --- | --- |
|  | PLV | awPLV |
| FPR | 0.037 (0.0296, 0.0462) | 0.0335 (0.0265, 0.0423) |
| FNR at 20% CNR | 0.774 (0.7552, 0.7918) | 0.156 (0.1408, 0.1726) |
| FNR at 30% CNR | 0.1805 (0.1643, 0.1980) | 0 (0, 0) |

### Empirical Results

Group ICA yielded 18 RSNs, as described in our prior report (3), shared on NeuroVault (10), and shown in Figure 3a. The subject’s gastric rhythm was normogastric, at 0.048 ± 0.001 Hz. Using FDR-corrected Wilcoxon rank tests, 11 of the 18 RSNs were found to be synchronized with the gastric BER; these results are shown in Figure 3c and tabulated in Table 2, which lists uncorrected *p*-values as well as *p*-values adjusted for multiple comparisons using the Benjamini-Hochberg method with an FDR of 0.05. (To facilitate comparison with earlier analysis, results based on PLV are shown in Figure 3b.)

**Fig 3.**
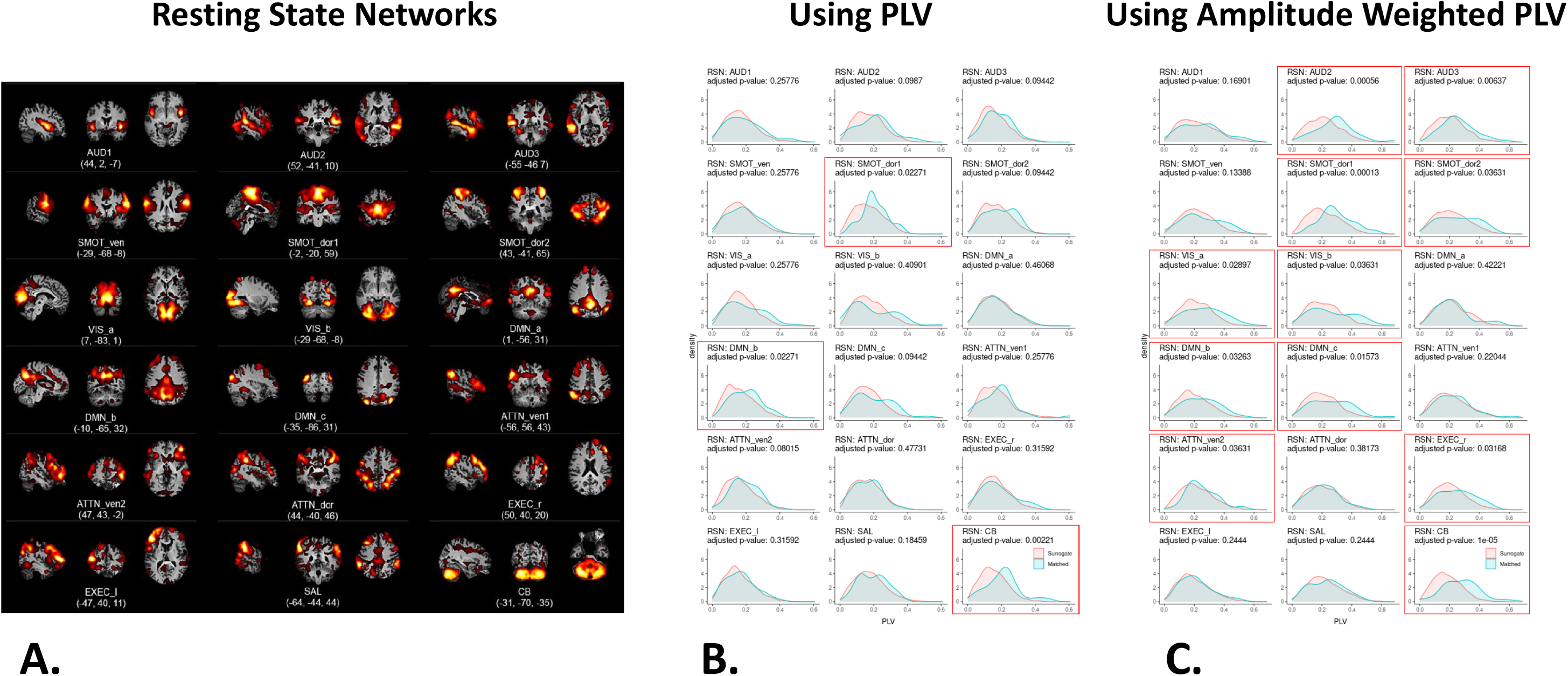
**A.** Aggregate spatial maps of resting-state networks (RSNs). Representative sagittal, coronal, and axial views (left-to-right) are overlaid on structural images in the Montreal Neurological Institute template space. Coordinates (in mm) for each view are given below each subfigure. (AUD: auditory, SMOT: somatosensory-motor, VIS: visual, DMN: default mode network, ATTN: attention, EXEC: executive, SAL: salience, CB: cerebellar, ven: ventral, dor: dorsal, r: right, 1: left). **B.** Gaussian kernel density estimates of PLV between resting-state networks and gastric signals recorded concurrently (cyan) and on different days (coral). *P*-values from Wilcoxon rank tests have been adjusted for multiple comparisons using an FDR of 0.05. Subfigures for networks with significant gastric synchronization are outlined in red. **C.** Gaussian kernel density estimates of awPLV between restingstate networks and gastric signals recorded concurrently (cyan) and on different days (coral). *P*-values from Wilcoxon rank tests have been adjusted for multiple comparisons using an FDR of 0.05. Subfigures for networks with significant gastric synchronization are outlined in red. (Parts A & B adapted from Choe, et al., 2021 (3).)

**Table 2.**
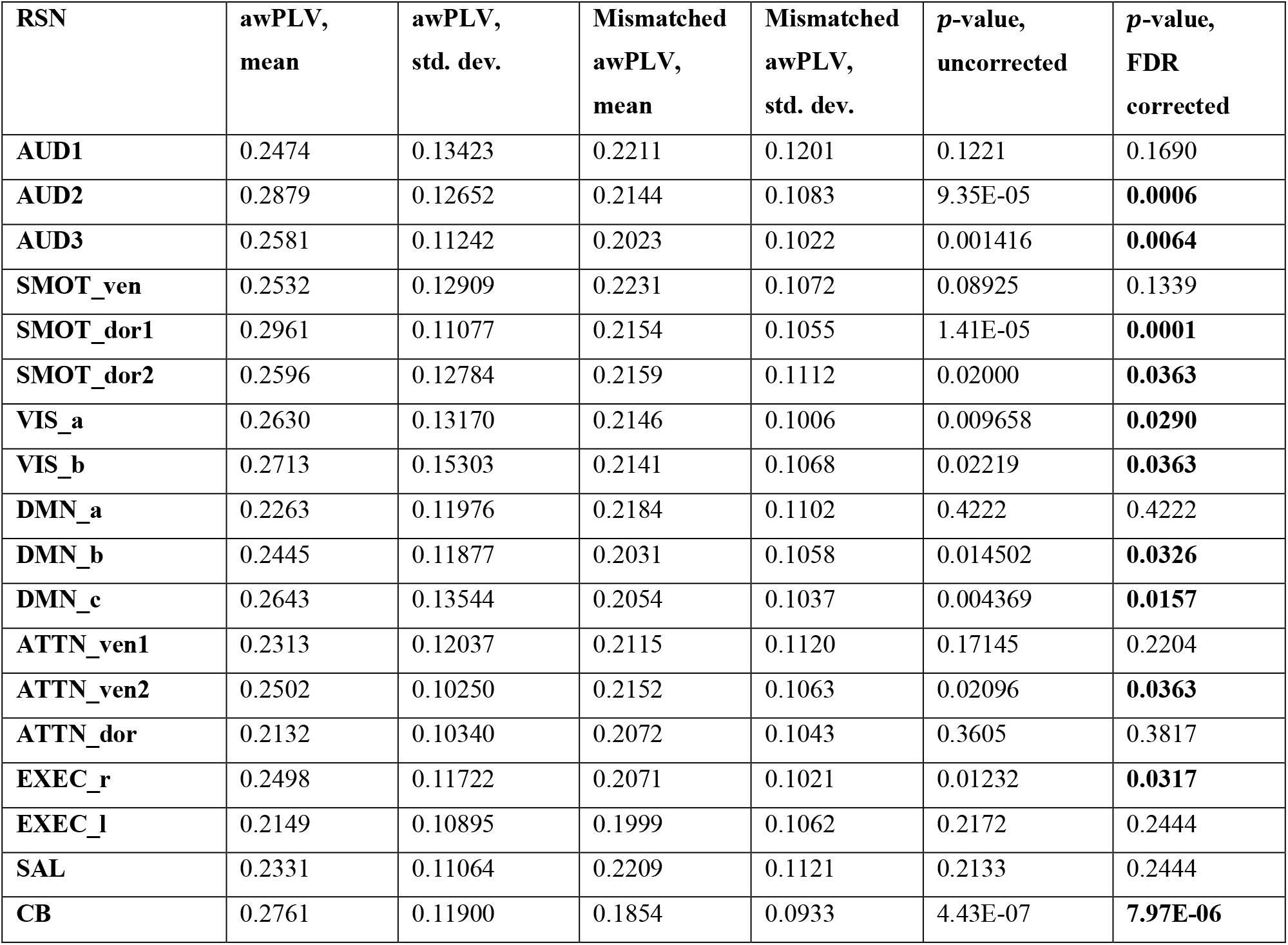
Amplitude weighted phase-locking value (awPLV) between resting-state networks (RSNs) and electrogastrography (EGG) signals. Corrected p-values of less than 0.05 are indicated in bold.

| RSN | awPLV,<br>mean | awPLV,<br>std. dev. | Mismatched<br>awPLV,<br>mean | Mismatched<br>awPLV,<br>std. dev. | p-value,<br>uncorrected | p-value,<br>FDR<br>corrected |
| --- | --- | --- | --- | --- | --- | --- |
| AUD1 | 0.2474 | 0.13423 | 0.2211 | 0.1201 | 0.1221 | 0.1690 |
| AUD2 | 0.2879 | 0.12652 | 0.2144 | 0.1083 | 9.35E-05 | **0.0006** |
| AUD3 | 0.2581 | 0.11242 | 0.2023 | 0.1022 | 0.001416 | **0.0064** |
| SMOT_ven | 0.2532 | 0.12909 | 0.2231 | 0.1072 | 0.08925 | 0.1339 |
| SMOT_dor1 | 0.2961 | 0.11077 | 0.2154 | 0.1055 | 1.41E-05 | **0.0001** |
| SMOT_dor2 | 0.2596 | 0.12784 | 0.2159 | 0.1112 | 0.02000 | **0.0363** |
| VIS_a | 0.2630 | 0.13170 | 0.2146 | 0.1006 | 0.009658 | **0.0290** |
| VIS_b | 0.2713 | 0.15303 | 0.2141 | 0.1068 | 0.02219 | **0.0363** |
| DMN_a | 0.2263 | 0.11976 | 0.2184 | 0.1102 | 0.4222 | 0.4222 |
| DMN_b | 0.2445 | 0.11877 | 0.2031 | 0.1058 | 0.014502 | **0.0326** |
| DMN_c | 0.2643 | 0.13544 | 0.2054 | 0.1037 | 0.004369 | **0.0157** |
| ATTN_ven1 | 0.2313 | 0.12037 | 0.2115 | 0.1120 | 0.17145 | 0.2204 |
| ATTN_ven2 | 0.2502 | 0.10250 | 0.2152 | 0.1063 | 0.02096 | **0.0363** |
| ATTN_dor | 0.2132 | 0.10340 | 0.2072 | 0.1043 | 0.3605 | 0.3817 |
| EXEC_r | 0.2498 | 0.11722 | 0.2071 | 0.1021 | 0.01232 | **0.0317** |
| EXEC_l | 0.2149 | 0.10895 | 0.1999 | 0.1062 | 0.2172 | 0.2444 |
| SAL | 0.2331 | 0.11064 | 0.2209 | 0.1121 | 0.2133 | 0.2444 |
| CB | 0.2761 | 0.11900 | 0.1854 | 0.0933 | 4.43E-07 | **7.97E-06** |

## DISCUSSION

We apply weighted phase-locking as a measure of synchronization of BOLD fMRI signals with the intrinsic electrical rhythm of the stomach. The awPLV was developed by Kovach (4) to improve sensitivity of phase-locking estimates (5, 6). The rationale is that when averaging phase-locking over a run, contributions should be weighted so as to reduce contributions from time points when the signal amplitude is small, and phase estimate is therefore noisy. In the present contribution, using the awPLV measure, 11 brain networks, which constitute a majority of the 18 networks estimated from the rsfMRI data, were found to be synchronized with the gastric rhythm. These 11 brain networks comprise three RSNs previously (3) found to be synchronized with the gastric rhythm using the non-weighted PLV measure—namely, a cerebellar network, a dorsal somatosensory-motor network, and a default mode network—as well as eight more: another dorsal sensory-motor network, two additional default mode networks, two visual networks, two auditory networks, a ventral attention network, and a right executive network.

The plasmode Monte Carlo simulation results indicate that the PLV measure can yield a high rate of false negatives (an FNR of 77%, under our modeled noise distribution). This suggests that differences seen in significance estimates based upon PLV vs. awPLV measures, in the respective analyses of our empirical data set, are likely due to false negatives using PLV. Neither PLV nor awPLV measures yield false alarm rates in excess of the specified 0.05 alpha rate.

A highly-sampled individual study design (11,12) was used. In order to test for statistical significance, an estimate of the chance-level distribution of gastric/RSN awPLV that would be seen in the absence of true stomach-brain phase synchronization was obtained using mismatched data (rsfMRI and EGG data acquired on different days). However, if the subject’s gastric frequency had been absolutely constant, then the gastric rhythm itself would be completely phase-locked across sessions and days. Hence, to the extent that the gastric rhythm is unexpectedly stable, our use of mismatched data as surrogate data would be overly conservative, and our estimates of significant synchronization would be understated. Relatedly, a limitation of this work is that it reports on data acquired in just one person; ongoing and future studies will ascertain whether the results reported herein generalize to a larger population.

Rebollo, et al., (13) were first to assess stomach/brain synchronization in concurrent rsfMRI and EGG data; they used the PLV because it tests for a consistent but arbitrary phase difference between signals, and because it was independent of signal amplitude. There are many other measures of synchronization available (14). The weighted PLV introduced by Kovach resembles the frequency-domain coherence measure (4). The application of synchronization measures to neuroelectrophysiological data has recently been reviewed (15).

Results reported herein imply communication between the enteric nervous system and several brain circuits not typically considered responsive to gastric state. Future work will interrogate hunger and satiety in healthy and disease states to probe these interactions and yield insight into their function. Three frameworks suggest that it may not be surprising to find activity in much of the brain that is partially but significantly synchronized with the stomach: First, interoception is vital for allostasis, which can be seen as a key driver of evolution of the large human brain; a large widely-distributed allostatic brain network has been reported (16). Second, it has recently been argued that all learning requires interoception (17), in which case, any mental activity that is influenced by or results from learning, in any way, should be seen as the product of an embodied, interoceptive, brain. Third, recent reports of widespread nervous system activity associated with a variety of ongoing behaviors (18) across a variety of organisms, would appear to be consistent with the present report of widespread brain activity associated with ongoing gastric electrical activity.

## CONCLUSION

This report introduces weighted phase-locking as a sensitive measure of synchronization of BOLD fMRI signals with the intrinsic electrical rhythm of the stomach. The *amplitude-weighted* Phase-Locking Value (awPLV) was used to assess synchronization between the gastric basal electrical rhythm and brain networks estimated from resting-state fMRI data using spatial independent component analysis. Using awPLV, a *majority* of resting-state brain networks were found to be significantly synchronized with the intrinsic rhythm of the stomach. Simulations suggest that differences seen in significance estimates based upon awPLV vs. PLV measures are likely due to false negatives using PLV. Empirical results imply communication between the enteric nervous system and brain circuits not typically considered responsive to gastric state.

## Footnotes

* Presented in preliminary form at the 2021 Annual Meeting of the International Society for Magnetic Resonance in Medicine.

